# Tuning molecular motor transport through cytoskeletal filament network organization

**DOI:** 10.1101/277947

**Authors:** Monika Scholz, Kimberly L. Weirich, Margaret L. Gardel, Aaron R. Dinner

## Abstract

The interaction of motor proteins with intracellular filaments is required for transport processes and force generation in cells. Within a cell, crosslinking proteins organize cytoskeletal filaments both temporally and spatially to create dynamic, and structurally diverse networks. The architecture of these networks changes both the mechanics as well as the transport dynamics; however, the effects on transport are less well understood. Here, we compare the transport dynamics of myosin II motor proteins moving on model cytoskeletal networks created by common crosslinking proteins. We observe that motor dynamics change predictably based on the microstructure of the underlying networks and discuss how this can be utilized by cells to achieve specific transport goals.

## I. INTRODUCTION

The interior of cells is organized by cytoskeletal structures. Protein monomers like actin and tubulin organize into polymers, resulting in polar filaments that can span cellular dimensions. These filaments interact with molecular motor proteins to drive intracellular transport, cellular shape change, and cell motility [1]. These different functions rely on different arrangements of the filaments, ranging from tight bundles in filopodia to meshes in the cell cortex [2].

Filaments are organized locally by proteins that can bind to multiple actin filaments to crosslink them into a network [2]. The type of crosslinking protein determines the resulting network structure. Proteins with a small crosslinking distance, such as fimbrin or *α*-actinin, create tightly packed bundles with the filaments spaced 8-36 nm apart [3, 4]. The crosslinking protein fascin creates similarly tight bundles, but the actin filaments are polarity-sorted as well [5, 6]. In contrast, filamin has a crosslinking distance of 160 nm, resulting in a loosely bundled structure for low concentrations of crosslinking protein [7]. While the consequences of these diverse structures for network mechanics have the focus of several studies [8–13], the consequences for motor transport have received less attention [14, 15].

We recently found that the microscopic arrangement of filaments can have profound effects on the transport of motor proteins across filament networks [16]. In particular, the presence of filament loops can lead to unproductive cycling of motors. These vortex-like states give rise to a power-law distribution of dwell times and glassy dynamics in intracellular transport. We showed that we could tune the transport properties of the motors by varying properties of the system that control the number of junctions that can support cycling. We thus predicted that cells could modulate their cytoskeletal structures spatially and temporally to control the motor dynamics.

Here, we demonstrate experimentally that actin filament network organization can modulate transport dynamics of myosin II motors. To isolate the effects associated with bundle architecture from those stemming from cellular regulation [17], we track myosin II on model networks of actin filaments bundled by biological crosslinking proteins. The dynamics can be interpreted in terms of the known features of the crosslinking proteins.

## II. TRANSPORT PROPERTIES DEPEND ON THE MICROSCOPIC BUNDLE ARCHITECTURE

We analyze single-particle tracking data from myosin II motors moving on a quasi-two-dimensional (quasi-2D) actin network (Fig. 1). The actin filaments are crosslinked with either *α*-actinin, fimbrin, fascin or filamin to create actin bundles with distinct microstructures (Fig. 1). The actin networks and skeletal-muscle myosin II were prepared as described in [16]. Single-particle trajectories were obtained using the Python-based implementation of the Crocker-Grier algorithm Trackpy [18]. The resulting trajectories align well with the crosslinked actin filaments (Fig. 1).

**FIG. 1.**
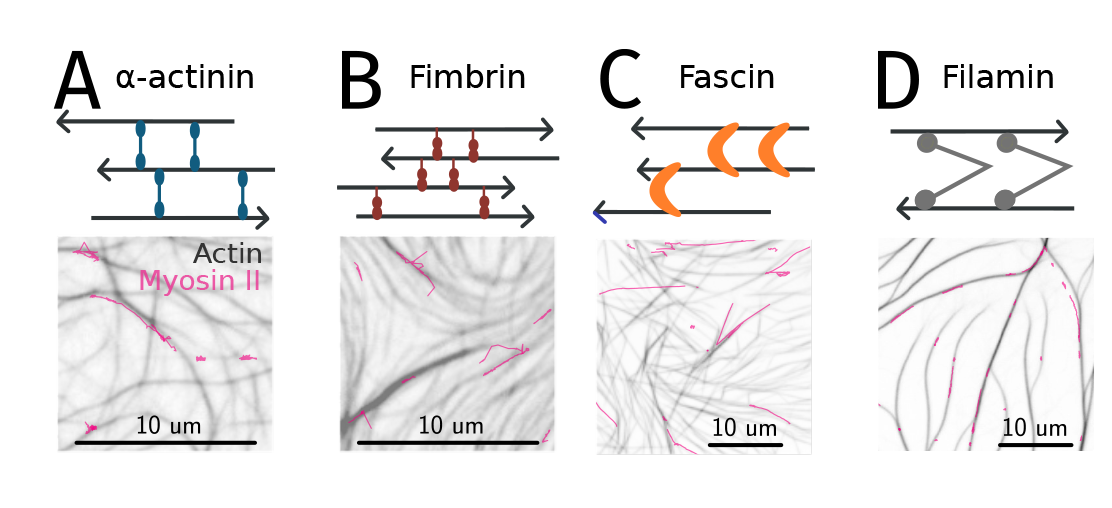
Schematic showing the interactions of four cross-linking proteins with actin filaments (red arrows). Fluorescence microscopy image of the quasi-2D actin networks after crosslinking with either *α*-actinin (A), fimbrin (B), fascin (C) or filamin (D). The trajectories of myosin II are overlaid (red). Note that the scale of the images change between panels, due to the large difference in myosin II speeds on different crosslinked actin networks.

For each experimental condition, we characterize the motor motion through the time-averaged mean-squared displacement (TA-MSD) as a function of lag time Δ and measurement time *T* as previously described [16, 19, 20]:

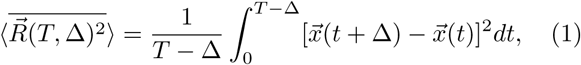

where 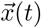 is the position of a motor at time *t*.

The exponent *α* in the scaling relation 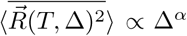 characterizes the motion: *α* = 1 for simple diffusion, and *α* = 2 for a purely inertial motion; non-integer values are possible as well (e.g., [19, 20] and references therein). We observe that the exponent of the TA-MSD as a function of lag time changes depending in the underlying bundle structure (Fig. 2). The polarity-sorted bundles created by the crosslinking protein fascin lead to an exponent close to two, indicating strongly directed motion. In contrast, the exponent of the TA-MSD is closer to one for filamin, *α*-actinin and fimbrin.

**FIG. 2.**
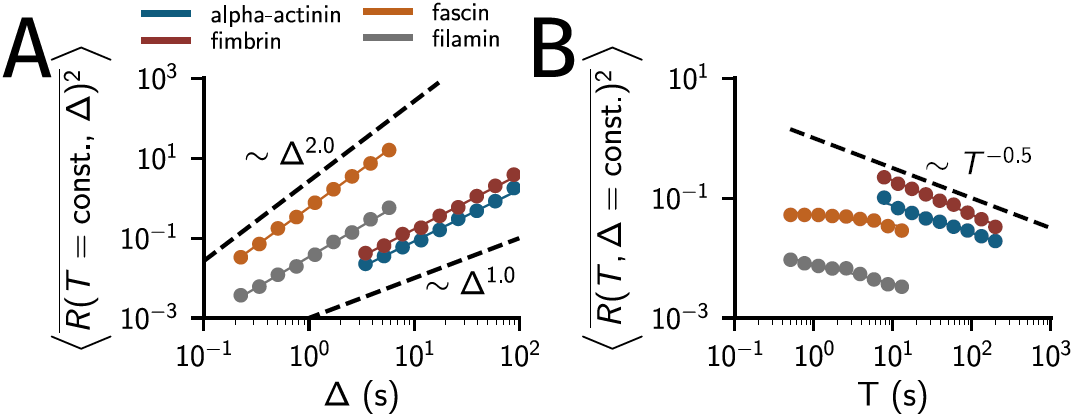
(A) Mean-squared displacement of myosin II minifilaments on actin networks crosslinked with different proteins as a function of lag time (Δ) and (B) as a function of measurement time (*T*). *T* = 9 s and Δ = 0.2 s for fascin (orange) and filamin (gray). For *α*-actinin (blue) and fimbrin (red), *T* = 137.7 s and Δ = 3.06 s. The number of trajectories is 210, 251, 236 and 256 for *α*-actinin, fimbrin, fascin and filamin, respectively. The mean trajectory lengths are 167, 208, 16 and 17 s for *α*-actinin, fimbrin, fascin and filamin, respectively.

We also consider the TA-MSD as a function of measurement time, *T*. This order parameter is sensitive to the ergodicity of the dynamics. If the properties of the transport process underlying the observed dynamics are unchanged over the course of measurement, the dynamics are ergodic, and the expected exponent is zero. However, as previously described for a fimbrin-crosslinked network, the mean-square displacement decreases as the measurement time increases (i.e., the exponent is less than zero) [16]. We find similar exponents for both *α*-actinin and fimbrin, which is unexpected considering the relevant scales. We previously described a mechanism for these non-ergodic dynamics that relies on motors interacting with at least two filaments at the same time [16] and getting effectively trapped. Filaments in fimbrin bundles are spaced closer than the binding radius of a myosin motor, and the myosin minifilament can interact with many filaments at the same time, supporting non-ergodic dynamics. However, the filament spacing in *α*-actinin bundles is larger than the myosin binding radius, which would suggest that the myosin minifilament spans these gaps via its long axis of 100–500 nm, which is considerably larger than the bundle spacing. The exponents for filamin and fascin do not deviate as strongly from zero, indicating that the dynamics are only weakly non-ergodic. In the case of fascin, the unipolar bundles favor directional transport, and do not support trapping, even if the motor protein interacts with multiple filaments at once. In the case of filamin the large filament spacing often leads to the motor protein interacting with a single filament.

## III. ANGULAR DISTRIBUTION ILLUSTRATES MICROSCOPIC DIRECTIONAL CHANGES

The TA-MSD cannot reveal directional correlations within trajectories, but there is evidence of such correlations in particle-tracking studies of molecular motors [19–24]. To quantify such correlations, we use the relative angle distribution [20]. The relative angle is defined by

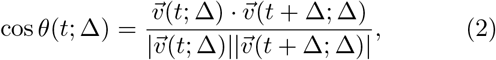

where 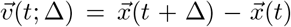. The normalized histogram (probability density function, PDF) of *θ* values for successive vectors within trajectories can be used as a directional order parameter.

The relative angle distribution is flat for simple diffusion because there are no correlations between steps of the random walk. A dictionary of relative angle distributions for a variety of more complicated transport processes can be found in [20]. Consistent with previous observations for molecular motors [20], for the four experimental conditions that we consider here (Fig. 3), we generally observe peaks at *θ* = 0 and *θ* = *π*, indicating an apparently inertial motion and frequent reversals, respectively.

**FIG. 3.**
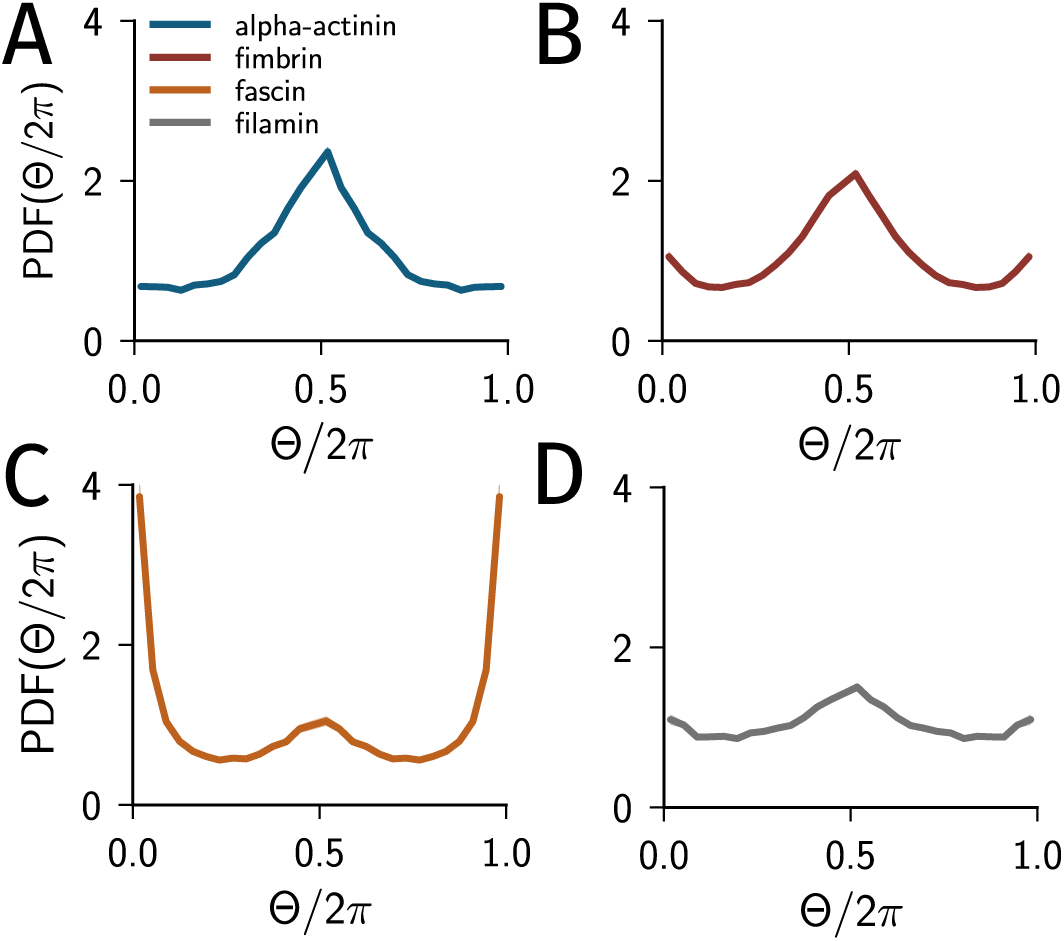
Relative angle distributions of myosin II motor trajectories on actin networks crosslinked with either *α*-actinin (A), fimbrin (B), fascin (C) or filamin (D). See text for discussion of the choice of Δ.

The relative angle distribution can be calculated at different Δ to elucidate the timescales contributing to the motion. To investigate the effects on the transport of local structure, as opposed to large-scale network topology, we chose a small Δ (Δ = 0.1, 0.1, 1.53, 1.53 s for fascin, filamin *α*-actinin and fimbrin, respectively). To compare recordings of the motion on different crosslinked networks, the magnitude of Δ was chosen to be inversely proportional to the mean motor velocity on a given network. For example, the mean velocity of myosin on the fimbrin-crosslinked network is only 38 nm/s whereas motors on the fascin-crosslinked network move at 620 nm/s. Thus, the respective values for Δ are 1.53 s and 0.1 s, respectively. We include all trajectories in the analysis, even those that are not moving significantly during our measurement. The average motor velocity on fascin bundles is smaller than the unloaded gliding speed of myosin on actin filaments [25, 26].

The motors on the fimbrin-crosslinked network show a strong peak at *π*, indicating that motors change direction frequently. This is consistent with tight, mixed polarity bundles that create an environment that supports tug-of-war [27, 28] and cycling [16] mechanisms. Again the results for *α*-actinin and fimbrin are similar, consistent with a trapping between the tighter bundles along the long axis of the myosin minifilament in both of these bundles. The relative angle distribution for *α*-actinin also agrees with a previously reported data set [20].

In contrast, the motor transport on fascin-crosslinked bundles shows a strong directional component, as evidenced by the peak at *θ* = 0 in the relative angle distribution. Since fascin arranges actin filaments into polarity sorted bundles, this is likely due to directed movement along a bundle. The loose bundles created by filamin result in a nearly flat angular distribution with only small peaks at *θ* = 0 and *π*. This indicates transport dominated by diffusion with only small directional components and few reversals.

Interestingly, the time-averaged mean-squared displacement shows a super-diffusive behavior for transport on the filamin-crosslinked network (Fig. 2A). Therefore, over large timescales the motor exhibits directed motion along the filamin network. However, on the smaller timescale used to calculate the angular distribution the motor shows nearly diffusional behavior. This could support two hypotheses. One is that the motors frequently hop between filaments in different microscopic orientations, such that the motion resembles a biased random walk. The other possible hypothesis is that the motors are detaching and diffusing within the bundles. Taken together, these measurements show that the relative angle distribution is sensitive to the bundle structure and can distinguish bundle structures resulting from different crosslinking proteins.

## IV. DISCUSSION

Using motor trajectories obtained from tracking myosin II motors moving on reconstitued actin networks, we find that transport depends on the microscopic structure of the filament network. Intuitively, polarity-sorted bundle structures lead to directional transport, whereas mixed polarity bundles result in a mixture of directed motion and trapped motion. Since the motor complex has a finite “reach”, the loose bundles formed by filamin are less effective at trapping the motor than the tightly spaced fimbrin bundles. From our results, one can deduce the dynamics on a hypothetical actin network with wide filament spacing but polarity-sorted bundles. Our results would suggest that the resulting motion would have a strong forward-directed component, but no significant trapping, thus resulting in apparently inertial dynamics over short timescales and simple diffusion over long timescales.

Our results have implications for the regulation of transport in cells, but is especially relevant for secreting cells, such as pancreatic beta cells. Although traditionally associated with microtubule-based transport, insulin secretory vesicles in beta cells have to cross the actin cortex before fusing with the membrane to release the hormone into the bloodstream [19]. The actin cortex is remodeled in response to insulin signaling [29]. It is likely that the timing of insulin release is affected by the structure of the actin cortex, and that the reported actin remodelling serves the purpose of making the network more amenable for traversing granules [30].

In summary, our results show that the organization of cytoskeletal networks has not only mechanical, but also dynamical consequences. The prevalence of different crosslinking proteins in different cellular regions, such as fillopodia, lamellopodia, or the cell cortex, could also be optimized for the transport processes that occur in those regions. Further studies should investigate the role of crosslinking proteins on motor dynamics *in vivo*. The principles that we elucidated could also be exploited to design novel materials with defined transport properties.

**TABLE 1.**
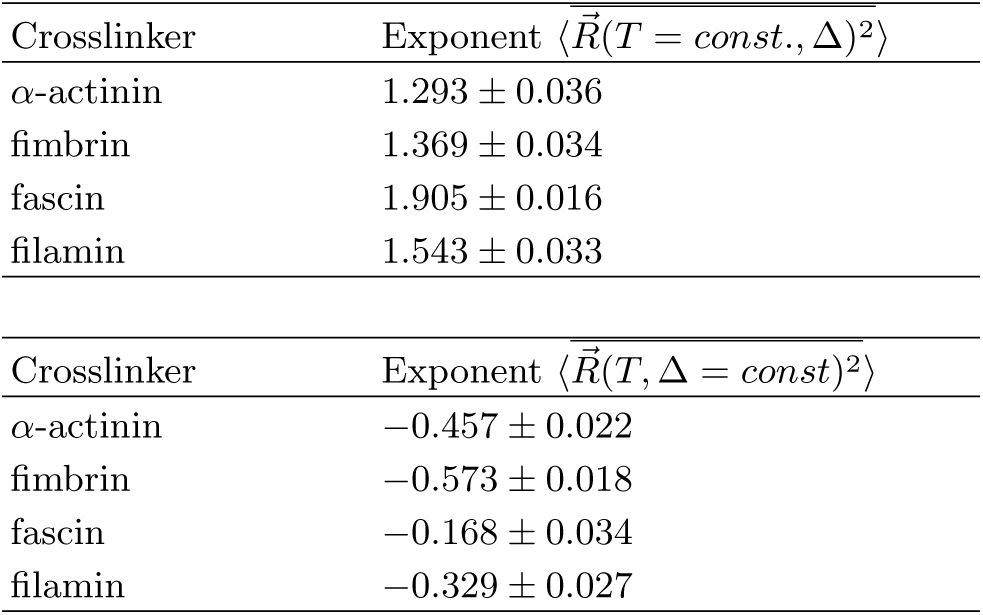
Exponents of the mean-squared displacement shown in Fig. 2. The error is the statistical error of the fit.

## V. ACKNOWLEDGMENTS

We thank Stanislav Burov for helpful discussions. We thank Samantha Stam, Todd Thoresen, and members of David Kovar’s laboratory (Jenna Christensen, Jonathon Winkelmann, and Youjii Li) for purified proteins. This work was primarily supported by the University of Chicago Materials Research Science and Engineering Center under award NSF DMR-1420709.

